# EC-PSI: Associating Enzyme Commission Numbers with Pfam Domains

**DOI:** 10.1101/022343

**Authors:** Seyed Ziaeddin Alborzi, Marie-Dominique Devignes, David W. Ritchie

## Abstract

With the growing number of protein structures in the protein data bank (PDB), there is a need to annotate these structures at the domain level in order to relate protein structure to protein function. Thanks to the SIFTS database, many PDB chains are now cross-referenced with Pfam domains and enzyme commission (EC) numbers. However, these annotations do not include any explicit relationship between individual Pfam domains and EC numbers. This article presents a novel statistical training-based method called EC-PSI that can automatically infer high confidence associations between EC numbers and Pfam domains directly from EC-chain associations from SIFTS and from EC-sequence associations from the SwissProt, and TrEMBL databases. By collecting and integrating these existing EC-chain/sequence annotations, our approach is able to infer a total of 8,329 direct EC-Pfam associations with an overall F-measure of 0.819 with respect to the manually curated InterPro database, which we treat here as a “gold standard” reference dataset. Thus, compared to the 1,493 EC-Pfam associations in InterPro, our approach provides a way to find over six times as many high quality EC-Pfam associations completely automatically.

## 1. Introduction

Proteins are macromolecules comprising one or more chains of amino acid residues. Protein molecules carry out many essential biological functions such as catalysing metabolic reactions and mediating signals between cells, for example. These functions are often carried out by distinct “domains”, which may often be identified as highly conserved regions within a multiple alignment of the sequences of a group of similar proteins, as in the Pfam database [1], for example. It is widely accepted that such protein domains often correspond to distinct and stable three-dimensional (3D) structures, and that there is often a close relationship between protein structure and protein function [2]. Indeed, it is well known that protein structures are often more highly conserved than protein sequences [3], and this suggests that proteins with similar structures will have similar biological functions [4]. The Protein Data Bank (PDB) [5, 6] now contains over 107,000 3D structures, most of which have been solved by X-ray crystallography or NMR spectroscopy. Structure-based classifications of protein domains such as SCOP [7] and CATH [8] have revealed many conserved structure-function relationships at the molecular level, and these classifications are now widely used in the community. However, because there does not exist a standard way to define a protein domain precisely, there is not always a one-to-one correspondence between domains defined by SCOP and those defined by CATH, for example, or between such structural domains and the domains defined by Pfam.

As well as sequence-based and structure-based classifications, proteins may also be classified according to their function. For example, the Enzyme Commission [9] uses a hierarchical four-digit numbering system to classify enzymatic function of many proteins. The first digit, or top-level “branch” of the hierarchy, selects one of six principal enzyme classes (oxidoreductase, transferase, hydrolase, lyase, isomerase, and ligase). The second digit defines a general enzyme class (chemical substrate type). The third digit defines a more specific enzyme-substrate class (e.g. to distinguish methyl transferase from formyl transferase), while the fourth digit, if present, defines a particular enzyme substrate. However, it should be noted that because EC numbers are assigned according to the reaction catalyzed, it is possible for distinct proteins to be assigned the same EC number even if they have no sequence similarity or if they belong to different structural families.

While the above classification schemes are very useful, they do not generally provide a direct relationship between enzymatic function and a 3D domain structure or a (sequence-based) Pfam domain. Thus, except for single-domain proteins where the mapping is obvious, unless a 3D structure has been very carefully annotated at the time it was deposited in the PDB (which is often not the case), it is generally not possible to compare and classify structure-function relationships at the domain level. Nonetheless, several groups have described approaches or resources that can associate PDB protein chains with enzyme EC numbers. For example, both the IMB Jena library [10] and the latest version of the PDBsum web site [11, 12] map each chain from a PDB file to its component CATH and Pfam domains, and each provides a link to the Enzyme database [13] for each PDB chain that has an EC number. PDBSprotEC [14] maps PDB chains to SwissProt and then uses the Enzyme database to obtain a mapping between SwissProt codes and EC numbers. Additional partial EC assignments are also retrieved directly from SwissProt. Columba [15] integrates annotation data from 12 different databases including PDB, SwissProt, CATH, SCOP, and Enzyme. For each PDB entry that has an EC number, Columba annotates the biological unit with the enzyme name and biochemical reaction, and it links SCOP and CATH domain information to each protein chain. PDB-UF [16] aims to assign EC numbers to unannotated protein structures which have no detectable sequence similarity to other proteins of known function. This approach first clusters existing protein structures using the 3D-hit structure alignment program [17]. It then assigns an unknown query structure to the most similar cluster, and it assigns a complete or partial EC number to the query using the EC numbers found in the cluster. Probably the most up-to-date and exhaustive association between PDB chains and EC numbers is currently provided by SIFTS [18], which is a collaboration between the Protein Data Bank in Europe and UniProt [19]. SIFTS incorporates a semi-automated procedure which links PDB chain entries to external biological resources such as Pfam, IntEnz [13], CATH and SCOP.

While all of the above approaches can provide associations between PDB protein chains and enzyme EC numbers, to our knowledge, SCOPEC [20] is the only published approach for automatically assigning EC numbers to structural domains. SCOPEC uses sequence information from SwissProt and PDB entries that have been previously annotated with EC numbers in order to assign EC numbers to SCOP domains. The SCOPEC approach first looks for PDB chains that fully map to SwissProt entries (to within up to 70 residues) and that match on at least the first three EC number digits. It then extracts the single domain structures which can thus be associated unambiguously with an EC number. It then uses these annotated domains as queries against the multi-domain structures to annotate homologous domains. It also uses the Catalytic Site Atlas [21] to locate catalytic domains in multi-domain structures. However, a limitation of the SCOPEC approach is that it normally associates EC numbers only with single domain proteins. Although SCOPEC can also propagate a known EC-domain association to a matching domain in a multi-domain protein, it is not designed to deconvolute EC-chain associations into individual EC-domain associations. Furthermore, it appears that the SCOPEC database is no longer available on-line. There is therefore a fresh need to develop a way of associating EC numbers with individual domains in order to study the large number of structural domains that now exist in the PDB.

Here, we present a novel statistical training-based approach for finding associations between EC numbers and Pfam domains directly from existing EC-chain associations from SIFTS and EC-sequence associations from SwissProt and TrEMBL. We call our approach “EC-PSI” (being short for “EC-Pfam statistical inferencing”). While SwissProt and TrEMBL were originally developed separately, both databases have since been incorporated in the UniProt resource. SwissProt is now a high quality, non-redundant, and manually curated part of UniProt Knowledge Base (UniProtKB). In contrast, TrEMBL is an automatically annotated and unreviewed section of UniProtKB, and contains around 40 times more entries than SwissProt. In order to parameterise and evaluate EC-PSI, we use the InterPro database [22] which contains a large number of manually curated Pfam-EC associations. Thus it may be used as a “gold standard” reference dataset against which our predicted Pfam-EC associations may be compared.

## 2. Methods

### a. Data Preparation

Flat data files of SIFTS (October 2014), SwissProt and TrEMBL (November 2014), and InterPro (version 48.0) were downloaded and parsed using in-house Python scripts. From the SIFTS data, we extracted associations between Pfam domains and PDB chains, and associations between PDB chains and EC numbers. These associations were imported into two tables of our relational database. In the first table, each PDB chain is related to one or more Pfam domains (FIG. 1A). In the second table, each PDB chain is related to one or more EC numbers (FIG. 1B). Thus, these two tables together define a many-to-many relation between EC numbers and Pfam domains (FIG. 1C). UniProtKB provides another source of relationships between EC numbers and Pfam domains. However, these relationships are mediated by UniProt accession numbers (ANs) instead of PDB chains. Since UniProtKB is divided into SwissProt and TrEMBL, we parsed and extracted the corresponding AN-Pfam and AN-EC associations from the SwissProt and TrEMBL databases, and we stored the resulting many-to-many relations in two further pairs of tables, similar to the two SIFTS tables.

**Figure 1.**
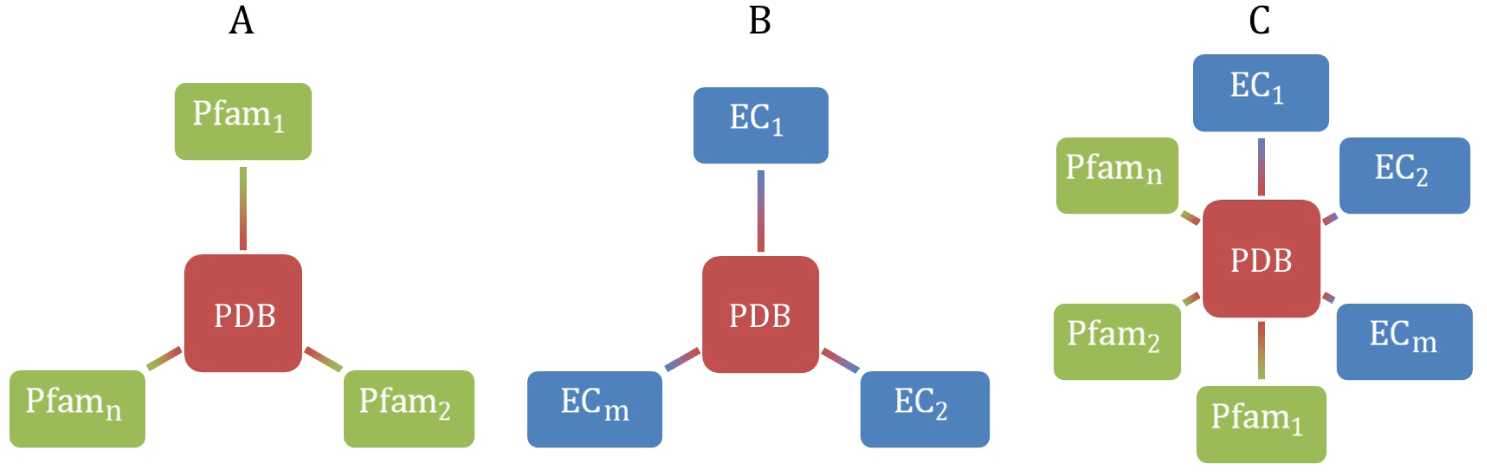
Illustration of the relationships extracted from the SIFTS database between (A) PDB chains and Pfam domains, (B) PDB chains and EC numbers, and (C) the many-to-many relationship between EC numbers and Pam domains.

As mentioned above, we used the InterPro manually curated EC-Pfam associations as a “gold standard” reference dataset. When considering only full four-digit EC numbers, we extracted a total of 1,493 EC-Pfam associations from InterPro, which we stored in our MySQL relational database. However, because we assume that all of the InterPro relations are “true” (i.e. correct) EC-Pfam associations, we needed to generate some plausible examples of false relations in order to train the EC-PSI algorithm. We therefore used our confidence score (see Section 2.b) to calculate and rank all possible EC-Pfam associations from SIFTS, SwissProt, and TrEMBL, and we extracted and stored 1,493 low-scoring EC-Pfam associations which could be calculated using data from at least two of the three databases. Because these associations have very little support in the data, we consider them to be “false” associations for the purpose of training our algorithm. In the rest of this paper, we will refer to the combined set of 1,493 “true” EC-Pfam associations from InterPro and our 1,493 calculated “false” associations as our “GoldStandard” dataset.

### b. Inferring Associations Between EC Numbers and Pfam Domains

In order to infer direct Pfam-EC relations from each of the above data sources, we collected all tuples of SIFTS data in the form (EC,PDB,Pfam), and we sorted these tuples by four-digit EC number and then by PDB chain in order to extract a tree-like set of relations for each EC number, as illustrated in FIG. 2. A similar sorting procedure was applied to the corresponding tuples extracted from the SwissProt and TrEMBL datasets. Then, for each EC number, we analyse its tree of associations by counting the numbers of occurrences of PDB chains (or ANs for SwissProt and TrEMBL) and Pfam domains. More specifically, for each Pfam domain within an EC tree, we calculate an EC-Pfam frequency score as the ratio between the number of chains in the tree that possess the given Pfam domain and the total number of PDB chains in the tree. In particular, letting *m* denote an EC number, *i* denote a PDB chain identifier, and supposing that the *m*^th^ EC tree contains *C*^*m*^ PDB chains denoted by 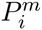(for *i* = 1,*…C*^*m*^) and that *D*_*n*_ denote the *n*^th^ Pfam domain, we define the PPFEC (“Pfam-PDB Frequency for a given EC-Pfam association”) score as

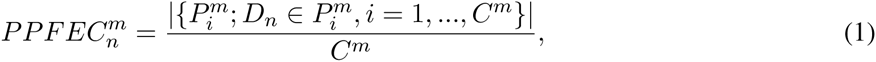

where |{*P* ^*m*^}| denotes the cardinality of a set of PDB chains. The notation 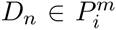 is understood to mean that chain 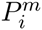 possesses domain *D*_*n*_. Equation (1) may be understood more graphically by considering FIG. 2. For a given EC number, *m*, and a given Pfam domain, *n*, the PPFEC is calculated as the degree of the Pfam node (number of connecting dashed lines) divided by the degree of the EC node (number of solid lines).

**Figure 2.**
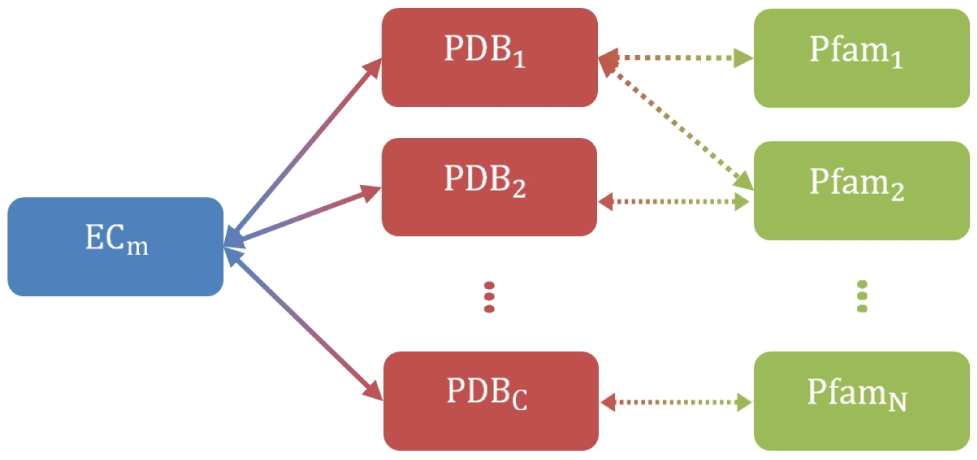
Graphical representation of the relationships between an EC number, *m*, and *N* Pfam domains via *C* PDB chains.

The corresponding frequencies for an inferred association between a Pfam domain and an EC number derived from the SwissProt and TrEMBL sequence annotations may be calculated in a similar way to give a “PSFEC” score (Pfam-SwissProt Frequency for a given EC-Pfam relation), and a “PTFEC” score (PfamTrEMBL Frequency for a given EC-Pfam relation), respectively. Thus, we obtain a frequency-based association score for each of the three data sources. However, because we wish to draw upon the relations from all three datasets, we combine the three frequency scores to give a single normalised “confidence score”,

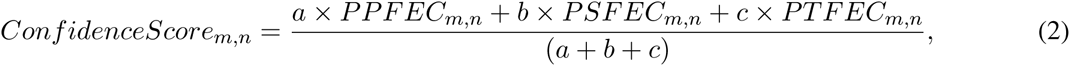

where *a*, *b*, and *c* are weight factors, to be determined, and where an individual frequency score is set to zero whenever there is missing data for a given *m* and *n*.

In order to find the best values for the above three weight factors, we varied their values from 0.0 to 1.0 in steps of 0.1, and for each combination we scored and ranked each of the 2,986 GoldStandard associations. Next, using the ranked list of true and false associations, we labeled true associations found in the top half of the ranked list as true positives (TPs), and we labeled true associations found in the bottom half of the list as false negatives (FNs). Similarly, we labeled false associations found in the top half of the list as false positives (FPs), and false associations in the bottom half as true negatives (TNs). We then calculated a Receiver-Operator (ROC) curve [23] of the TP rate against the FP rate, and we used the area under the curve (AUC) of the ROC plot as the overall quality measure of the scoring function.

### c. Defining a Confidence Score Threshold

Given that the best weights for each data source have been determined, we next wished to determine an overall threshold for our EC-Pfam association confidence score. In order to do this in an objective way, we randomly split the GoldStandard dataset into two equal groups with equal numbers of true and false instances to give a “Training” dataset and a “Test” dataset. Next, we scored and ranked the members of the Training dataset, and we divided the ranked list into two subsets according to a threshold value that ranged from 0.0 to 1.0 in steps of 0.01. For each threshold value, we counted the number of TPs (true associations above the threshold), FPs (false associations above the threshold), TNs (false associations below the threshold), and FNs (true associations below the threshold). We then calculated the recall, *R*, precision, *P*, and their harmonic mean in order to obtain the “F-measure” according to

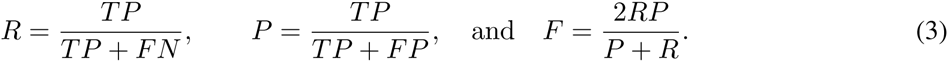

The score threshold that gave the best F-measure was selected as the best threshold to use for accepting predicted associations.

## 3. Results and Discussion

### a. Parameters of Our EC-PSI Procedure

Our EC-PSI procedure takes as input three large datasets of EC-chain associations from SIFTS, and EC-sequence associations from SwissProt and TrEMBL. These individual source datasets, which contain 6, 204, 9, 879, and 28, 572 associations respectively, were merged to give a global dataset of 32,018 non-redundant EC-Pfam associations. Our scoring function was trained using our GoldStandard dataset consisting of 1,493 “true” associations taken from InterPro and 1,493 “false” associations taken from low-scoring associations from SIFTS, SwissProt, and TrEMBL. We found that the best ROC-plot AUC is obtained with weights *a* = 0.1, *b* = 1.0, and *c* = 0.1 (Section 2.a), for a maximum AUC value of 0.888. These weights clearly give a 10-fold greater importance to the associations derived from SwissProt than to those derived from SIFTS and TrEMBL.

Using these weights, various threshold values of the confidence score were tested on the “Training” subset of our GoldStandard dataset, using the F-measure to quantify the results objectively (Section 2.b). The optimal score threshold was found to be 0.08 for a maximum F-Measure of 0.828. Applying this threshold to the GoldStandard Test subset yielded a comparable F-measure value of 0.810, and precision and recall values of 0.948 and 0.707, respectively. This threshold was then used to infer new EC-Pfam relations from the merged dataset, with a confidence score for each association being calculated by our scoring function.

### b. Global Analysis of Calculated EC-Pfam Associations

The results of the filtering process are summarized in Table 1. This table shows the numbers of EC-Pfam associations along with the numbers of distinct EC numbers and Pfam entries involved in those associations for the three source datasets, our merged global dataset before and after filtering (the latter corresponding to our “calculated” EC-Pfam associations), and for the InterPro dataset of true associations. The overlap between these two last datasets is shown in the last line of the table.

**Table 1.** Statistics on the given and calculated EC-Pfam associations.

| Dataset | EC-Pfam associations | 4-digit EC numbers | Pfam entries |
| --- | --- | --- | --- |
| SIFTS | 6,204 | 2,575 | 2,606 |
| SwissProt | 9,879 | 3,959 | 3,147 |
| TrEMBL | 28,572 | 3,538 | 5,839 |
| Merged | 32,018 | 4,588 | 6,290 |
| InterPro | 1,493 | 676 | 1,273 |
| EC-PSI (calculated) | **8,329** | **4,436** | **2,462** |
| Common to EC-PSI and InterPro | 1,089 | 592 | 944 |

Overall, Table 1 shows that our EC-PSI procedure yielded a total of 8, 329 calculated EC-Pfam associations that include 1, 089 associations already present in InterPro. While this shows that EC-PSI finds 73% (100 *\** 1, 089/1, 493) of the “correct” EC-Pfam associations in InterPro, it also shows that 27% (404/1, 493) of correct InterPro associations have EC-PSI confidence scores below our chosen score threshold of 0.08. This relatively high proportion of “missed” associations reflects the fact that our EC-PSI method is designed to discover EC-Pfam associations with strong factual support, whereas InterPro contains a large number of low frequency expert-annotated associations. More specifically, the score threshold of 0.08 was chosen to give a good tradeoff between precision and recall through the F-score. If, for example, the score threshold is reduced from 0.08 to 0.01, the recovery of correct InterPro associations increases to 90% (1,354/1,493), but the number of “false” InterPro associations rises from just 75 to 822.

Given that InterPro may be considered to represent the largest manually curated source of Pfam-EC associations currently available, it is interesting to consider the relative increase in the number of associations that our EC-PSI approach can provide. We therefore calculated as ratios (or “scale-up factors”) the differences between the associations calculated by EC-PSI and those of InterPro in terms of the total number of associations and the numbers of distinct EC numbers and Pfam entries involved in those associations. In FIG. 3, the scale-up factors are displayed across the 6 top-level branches of the EC classification (1-6) and for the entire datasets (All). It can be seen that the scale-up factors for EC-Pfam associations and for EC entries reach their maximum levels in branch 1 (oxydoreductases), and that they fluctuate around their average values (All) rather evenly in all other branches, with branch 6 (ligases) having the lowest values. The same is true for the scale-up factor for the number of Pfam entries, but the difference is less marked in branch 1. In fact, the average increase in Pfam entries is only about 2-fold compared to about 6-fold for Pfam-EC associations and EC numbers. This is consistent with the fact that not all Pfam entries can be assigned an EC number because not all Pfam domains are associated with an enzymatic activity.

**Figure 3.**
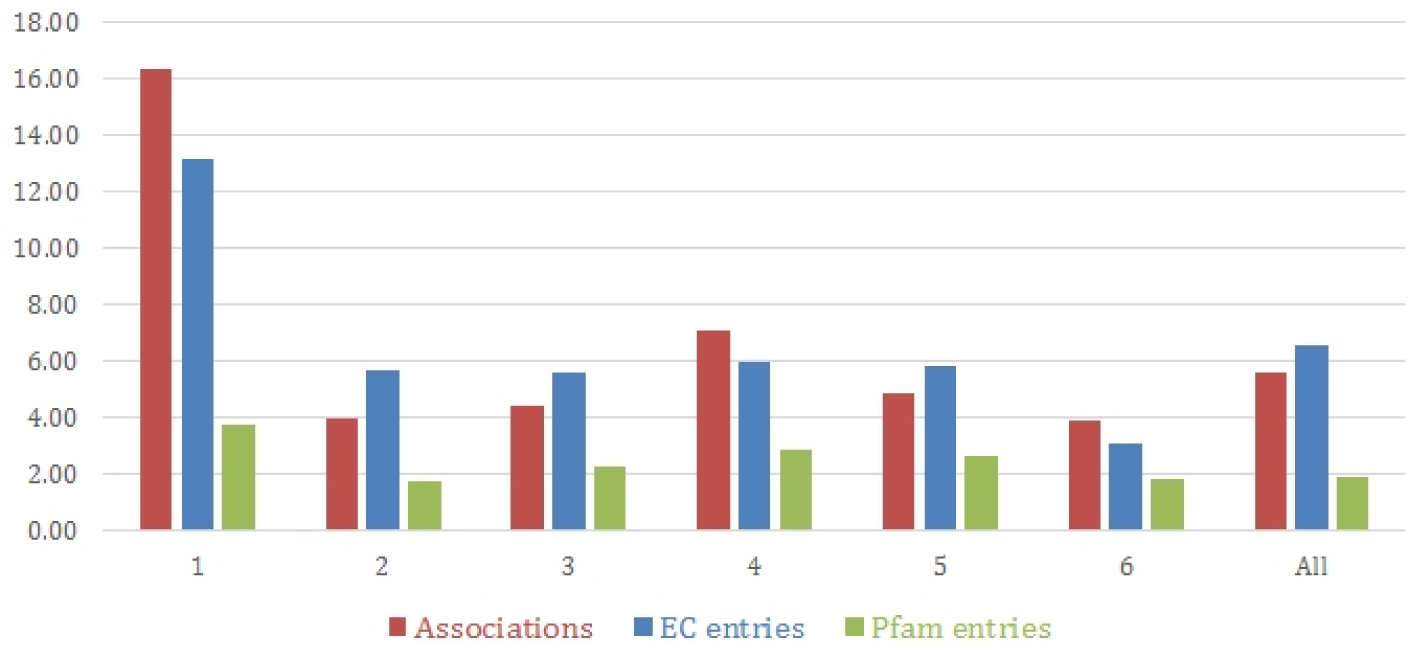
Scale-up factors for the EC-PSI and InterPro associations according to the EC branch. 1 : oxydoreductases ; 2 : transferases ; 3 : hydrolases ; 4 : lyases ; 5 : isomerases ; 6 : ligases ; All : all EC numbers.

### c. Comparison Between Calculated and InterPro EC-Pfam Associations

In FIG. 4A, the average number of EC-Pfam associations is plotted per EC number (1) and per Pfam entry (2) for both InterPro and our calculated dataset. The ratios are very close for the EC numbers (2.2 and 1.9, respectively), suggesting that our method follows the quality of annotation of InterPro and does not propose an excess of possibly incorrect EC-Pfam associations. On the other hand, the ratio is much higher for Pfam entries (3.38 versus 1.17), which reflects a significant enrichment in the annotation of Pfam domains. The rest of the figure shows the distribution of EC numbers (B) and Pfam entries (C) with respect to the number of associations they are involved in. Clearly the proportion of EC numbers (respectively, Pfam entries) that are involved in only one EC-Pfam association is reduced in our calculated dataset.

**Figure 4.**
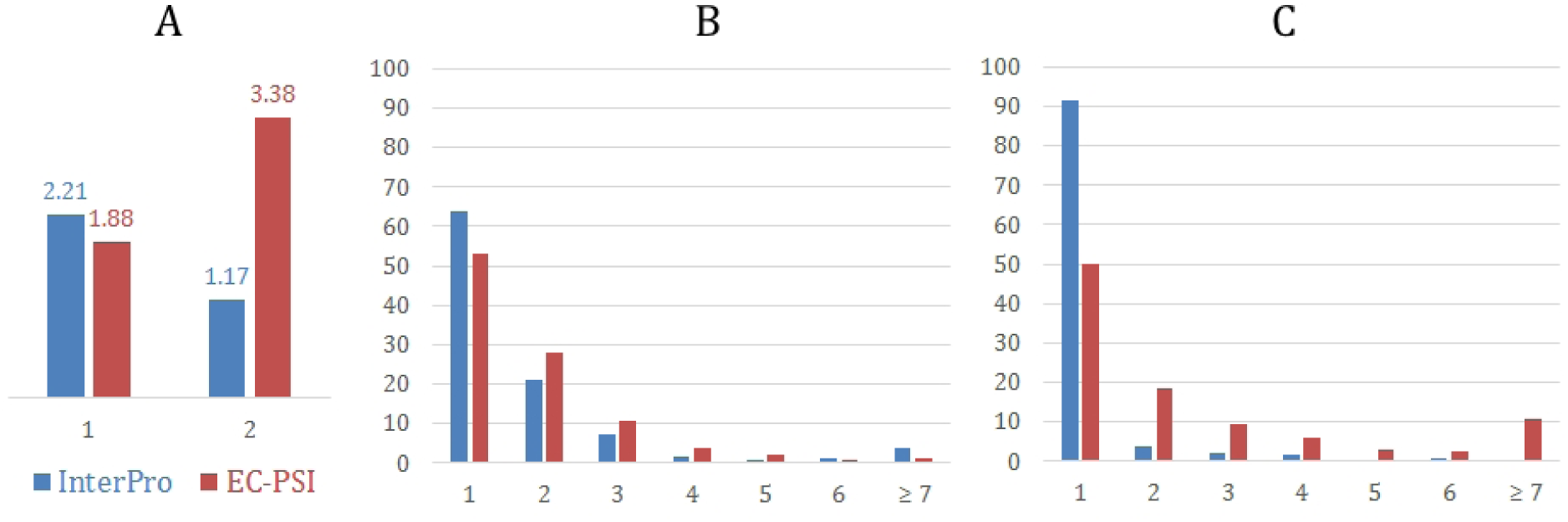
A : average number of EC-Pfam associations per EC number (1) and per Pfam entry (2) for the InterPro (blue) and calculated EC-PSI (red) datasets. B : distribution of EC numbers according to their numbers of associations with Pfam entries. C : distribution of Pfam entries according to their numbers of associations with EC numbers.

Overall, FIG. 4 shows that our collection of EC-Pfam associations rather favours multiple associations, thereby reflecting the complex many-to-many relationships that exists within the original datasets. Furthermore, many of the multiple associations calculated by EC-PSI seem to be quite reasonable from a biological point of view. For example, EC-PSI finds the unique InterPro association between EC 6.1.1.9 (valine-tRNA ligase) and the Pfam domain PF10458 (Valyl tRNA synthetase, tRNA binding arm) with a confidence score of 0.781, but it also finds two further associations with the same EC number that are not in InterPro, namely with PF08264 (tRNA anticodon-binding domain ; EC-PSI score 0.976) and PF00133 (tRNA synthetases class I ; ECPSI score 0.997). These two additional associations complete the biological picture of a tRNA ligase because they comprise a second constitutive domain, in addition to PF10458, of this complex enzyme. On the Pfam side, another interesting example is the unique InterPro association between PF04715 (Anthranilate synthase component, N terminal region) and EC 4.1.3.27 (anthranilate synthase). EC-PSI finds this association with a score of 0.522, but in addition it finds two further associations for the same Pfam domain, namely with EC 2.6.1.85 (aminodeoxychorismate synthase, EC-PSI score 0.675) and EC 2.6.1.86 (2-amino-4-deoxychorismate synthase ; EC-PSI score 0.833). In this case, the multiple association found by EC-PSI for PF04715 may be explained by the fact that all three enzymes share a common substrate (i.e. chorismate).

### d. Future Work

The approach presented here calculates associations using four-digit EC numbers. However, because EC numbers have an embedded hierarchy, and because it seems reasonable to suppose that enzymes that act on similar substrates are likely to be evolutionarily related, it could be interesting to consider making additional associations by collecting and analysing less specific three-digit associations. This could provide a way to infer additional associations that have weak direct support (low four-digit confidence scores), but which have good support at the three-digit level. We plan to analyse the support of EC numbers associated with more than one Pfam entry in order to detect those EC numbers that correspond to combinations of domains (in other words to detect cases where two or more domains are physically necessary to support a given enzyme function). We also want to improve the way that candidate associations from different sources are combined. Even though our current scoring function gives 10 times more weight to SwissProt than SIFTS and TrEMBL, it is still useful use all three data sources because our algorithm finds 312 EC-Pfam associations from SIFTS and 797 from TreMBL which are not present in the SwissProt data. However, it would be desirable to use a more statistically sound measure of the reliability of each data source, perhaps based on a comparision of the associations found after random shuffling of the data, for example.

## 4. Conclusions

Given the extensive protein chain/sequence annotations that now exist in the SIFTS, SwissProt, and TrEMBL databases, there is a need to be able to exploit this rich knowledge at the protein domain level. We achieved this aim by first collecting existing associations between EC numbers and protein chains or sequences, and then by using a statistical training-based scoring method to analyse the many-to-many relations embedded in these data. Using the above data sources, our approach is able to infer a total of 8,329 direct EC-Pfam associations. Thus, compared to the 1,493 manually curated InterPro EC-Pfam associations, our approach provides a way to find over six times as many associations completely automatically. We have also proposed some possible ways to extend and further analyse the coverage of the EC-PSI approach. We believe that the large numbers of EC-Pfam associations calculated using our approach can contribute considerably to enriching the annotations of PDB protein chains, and that this will facilitate a better understanding and exploitation of structure-function relationships at the protein domain level.

## Acknowledgements

This project is funded by the Agence Nationale de la Recherche (grant reference ANR-11-MONU-006-02), the Institut National de Recherche en Informatique et Automatique (Inria), and the Lorraine Region.

